# Nonlinear integration of sensory and motor inputs by a single neuron in *C. elegans*

**DOI:** 10.1101/2025.04.05.647390

**Authors:** Amanda Ray, Andrew Gordus

## Abstract

Context is important for sensory integration. Rather than simply considering sensory information independently, the brain integrates this information to inform behavior, however identifying this property at the single-neuron level is not trivial. In *Caenorhabditis elegans*, the paired interneurons AIBL and AIBR (AIB) have a compartmentalized organization of presynapses along its singular process. Sensory and sensory interneurons primarily synapse along the proximal process, while motor and motor interneurons synapse along the distal process. Since this neuron has graded potentials, the simplest model for AIB integration is simply a convolution of its presynaptic inputs. Through a series of experiments to manipulate sensory and motor input onto AIB, we find that while AIB activity is primarily a convolution of motor inputs, its sensory responses are not integrated independently. Instead, the gain in sensory input is a function of the temporal dynamics of motor input. Sensory information is reinforced when it matches the expected behavioral response. We find this property is also observed in other whole-brain datasets. Context-dependent behavioral responses to sensory input is well-documented. Here, we show this property can be localized to single neurons in the worm nervous system. This integration property likely plays an important role in context-dependent decision-making, as well as the highly variable dynamics of the worm nervous system.

## Introduction

The role of each neuron in the brain is to integrate different sources of presynaptic and extra-synaptic inputs to influence synaptic release. In mammalian neurons, dendritic branches serve as individual computational units. Previous reports have proposed a two-layer model of integration where each dendrite processes inputs separately and are then summed collectively to the cell as a whole ^1,2^. With a stereotyped connectome that was recently updated, the nematode *Caenorhabditis elegans* offers the opportunity to reliably stimulate and record from the same pre and postsynaptic pair of neurons reliably across different individuals ^3^ ^4^. Most worm neurons have graded potentials that lack dendritic arbors, and instead have processes in which pre and post-synaptic contacts occur *en passant* ^3,4^. While many of these synapses appear randomly distributed, several interneurons possess synapses along their processes which are bunched, which may enable local or nonlinear computation that gets integrated locally by the postsynaptic neuron ^5^.

One of the neurons in the worm connectome that exhibits synaptic bunching is AIB, an interneuron that integrates sensory and motor-driven inputs, and drives motor behavior ^3,6–9^ Along the AIB process, there’s a unique bias between these sensory-related and motor-related synapses, unlike other worm neurons which tend to have more randomly arranged presynaptic partners along the process ^3,4^. Chemosensory neurons and sensory interneurons tend to synapse along the proximal side of AIB’s process closest to the cell body, while neurons associated with motor behaviors lie along the opposite distal end of AIB’s process (Figure 1A, Table 1) ^3,4^.

**Figure 1:**
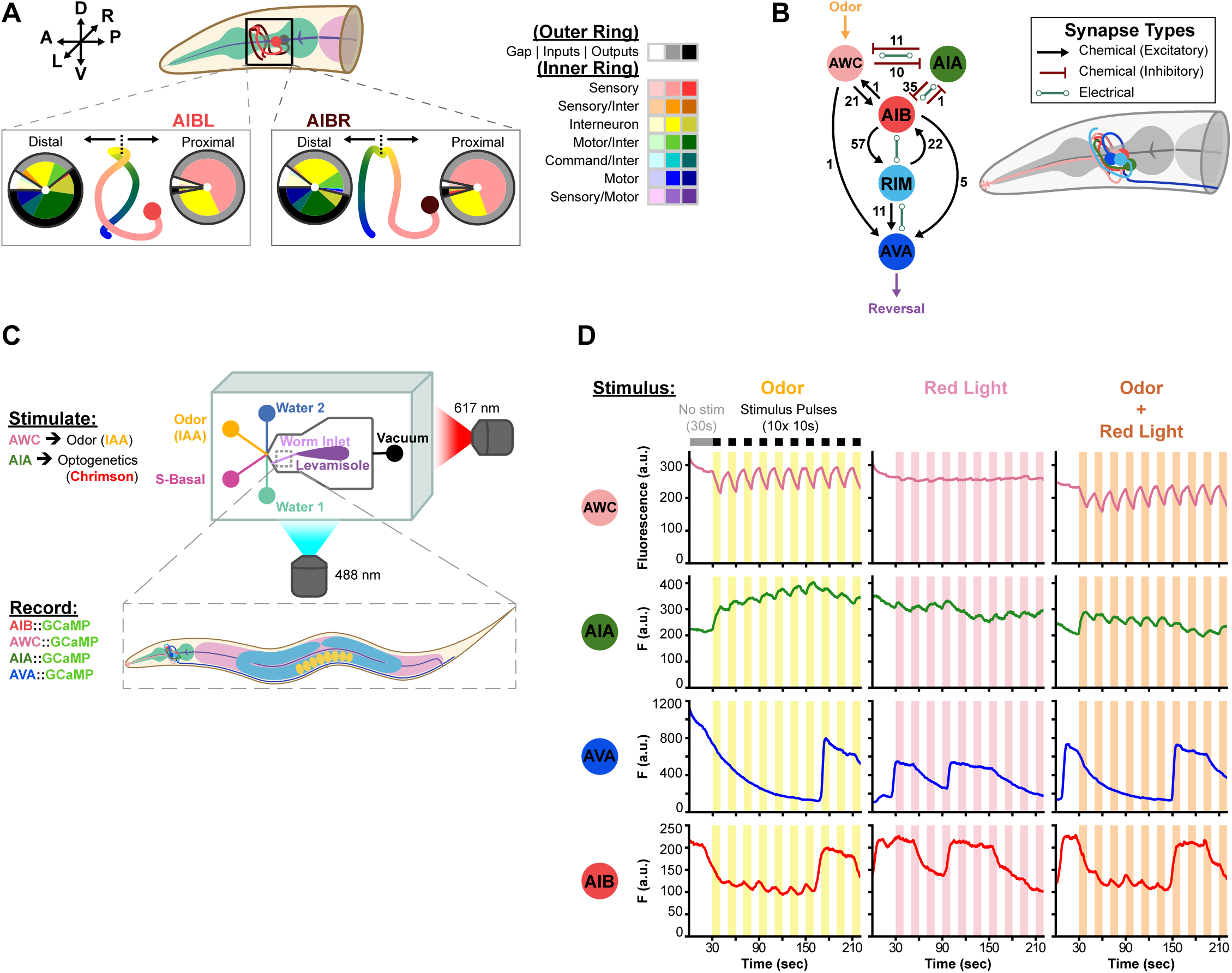
The synaptic partners of AIB. **A)** (Top) Diagram of AIBL (red) and AIBR (dark red) in the head of *C. elegans*. (Below) AIBL/R color-coded based on the types of synaptic partners that lie along the proximal and distal ends of the AIB process. Synaptic partner compositions are shown in nested pie charts beside the cartoon neurons; the outer rings denote type of synapse (gap in white, inputs in gray, and outputs in black), the colors of the inner ring correspond to the function of each synapse (sensory, interneuron, motor, command, and various polymodal combinations) and the shades within each color correlate with the type of synapse (i.e. light blue is a gap junction between AIB and a motor neuron, blue is presynaptic motor neurons onto AIB, and dark blue is for motor neurons postsynaptic to AIB). *See Supplementary Table 1 of synaptic composition, data obtained from the worm wiring database (*https://wormwiring.org/apps/neuronVolume/) ^4^. **B)** Selection of AIB synaptic partners with number and types of synapses. Cartoon on the right illustrates relative locations of soma and processes of each neuron in the head of the worm (only left side partners of each neuron are shown for simplification). **C)** Graphic of experimental setup to perform live calcium imaging of worms restrained in chip. Isoamyl alcohol (IAA) was delivered during imaging to stimulate AWC, and an external red LED stimulated AIA via Chrimson. **D)** GCaMP fluorescence of AWC, AIA, AVA, and AIB in response to stimuli. Example trace from a single AWC::GCaMP7s worm and a single AIB-AIA-AVA GCaMP7s worm across one full experiment (AIA, AIB, and AVA recordings were performed together but are separated out here for clarity). Phases were recorded within a minute of each other across the same worm for one full experiment (note the continuity of each trace between phases). Recordings were initialized with 30 seconds of no stimulus, followed by 10 bouts of 10 second ON/OFF stimuli pulses.

Despite integrating both sensory and motor information, AIB activity most closely correlates with the worm reverse-motor program ^7,9–12^. Most of AIB’s postsynaptic partners are motor-related neurons, and when optogenetically activated, AIB increases the likelihood of reversal behavior ^13,14^. Conversely, optogenetic activation of motor neurons presynaptic to AIB also drive AIB activity ^9^, which indicates AIB participates in a reinforcing feedback loop for reversals. However, the strength of sensory drive onto AIB is strongly influenced by competing input from motor activity from RIM, and in the absence of motor activity, AIB is more likely to produce activity that correlates with sensory input ^9^. How AIB integrates these competing inputs is not well-established.

To investigate the integration properties of AIB, we chose 4 synaptic partners of AIB: AWC, AIA, RIM, and AVA (Figure 1B). AWC is an excitatory, well-studied chemosensory neuron that can be activated by isoamyl alcohol ^8,15,16^. AWC can variably drive activity in AIB based on odor stimulation ^9^. AIA is an important pre-synaptic partner of AIB and accounts for roughly 12% of the presynaptic input out of a total of 288 inputs from 50 neurons presynaptic to AIB ^4^. AIA in turn integrates inputs from other sensory neurons like AWC ^17,18^. The RIM interneuron is strongly coupled to the AVA command neuron to generate reversal locomotion ^7,9,19–21^, and is the primary postsynaptic partner of AIB with ∼37% of the synapses out of a total of 155 outputs from 43 neurons postsynaptic to AIB ^4^. This combination of candidate neurons provides a simple sensorimotor network to study integration, where AWC and AIA provide sensory information onto AIB, and RIM provides motor information.

The simplest model of AIB integration is that it integrates presynaptic inputs linearly and independently. For example, when AVA is optogenetically stimulated, it can induce a graded potential in AIB that appears to be a convolution of AVA activity ^9^. However, AIB responses to sensory input is highly variable ^9,10^. This variability is observed behaviorally as well; even though worms may have a preference for a certain odor, individual behavioral responses to odor are highly volatile between individuals, and within an individual over multiple trials ^9,10^. Behavioral responses to sensory stimuli can be highly contextual. How reliably a worm responds to touch stimuli is strongly dependent on its behavioral state ^22,23^. This implies that presynaptic inputs may not necessarily be independent, and postsynaptic processing may rely on cross-dependencies between presynaptic or neuromodulatory partners. Here we examine the integration properties of AIB, and how well nonlinear integration assumptions reliably explain AIB responses to presynaptic activity.

## Results

### A combination of sensory and motor synaptic partners drive AIB activity

AIB is a first layer amphid interneuron that processes information between many sensory and motor neurons and other interneurons ^3,4^. A detailed look at the composition of all the synaptic partners of AIB and their locations along its process ^4^ reveals discrete patterning of presynaptic neuron identities (Supplementary Table 1). Most of the proximal presynaptic inputs are from amphid sensory neurons, and most of the distal synapses are comprised of presynaptic and postsynaptic motor-related neurons (Figure 1A). Among the interneuron synaptic partners, many of them are like AIB and integrate sensorimotor information, including first layer amphid interneurons (AIA, AIY, AIZ). This polarized separation of presynaptic identities showcases AIB’s role in synthesizing sensory information to drive motor behavior.

A subset of presynaptic neurons was chosen to investigate the sensorimotor integration properties of AIB (Figure 1B). AWC is an amphid chemosensory neuron that detects odorants ^8,15^, and is known to drive AIB activity ^8,9^. AIA is AIB’s primary presynaptic partner on the proximal process. AIA is a sensory interneuron that receives many direct inputs from chemosensory neurons, including AWC ^17,18^. AIA has several inhibitory synapses onto AIB and AWC, and its activity is typically anti-correlated with AIB and AWC ^6,18^. AIB shares several pre and post synapses with the RIM interneuron, which in turn synapses onto the AVA command neuron to drive reversal behaviors, and is tightly coupled to AVA activity ^9,11,12^.

To monitor the activity of AIB in the context of sensorimotor inputs, we expressed the genetically encoded calcium indicator GCaMP7s ^24^ in AIB, AIA, and AVA together in one strain (Figure 1C). Since RIM is in close proximity to AIB, we recorded from AVA as a proxy for RIM activity, as they have very closely correlated activity ^9,11,12^. Since the soma of AIA has very dampened activity compared to the neurite ^18^, we quantified neurite fluorescence in the same plane as the AVA and AIB somata. The soma from AWC is more lateral than the somata from AVA and AIB, and was recorded separately. AWC has a robust and consistent hyperpolarization in response to the odorant isoamyl alcohol, IAA, and repolarizes upon removal of odor ^8,9,16^, while the responses of AIB, AVA, and AIA to odor were more variable (Figure 1D). We also expressed the red-shifted channelrhodopsin, Chrimson ^25^, in AIA in our triple-GCaMP strain to optogenetically stimulate AIA in the presence or absence of odor pulses. During our recordings, AVA is frequently sporadically active, allowing us to monitor spontaneous motor activity without a need for direct stimulation. Thus, we were able to monitor and stimulate AWC and AIA with odor and optogenetics, respectively, to deliver sensory inputs to AIB, while simultaneously monitoring motor activity from AVA. Together, this allowed us to observe AIB activity in the context of varying sensory and motor inputs.

To stimulate AWC and AIA individually and simultaneously, the delivery of sensory inputs (odor and red light) was separated into three different phases for each worm – IAA odor only, red light only, and odor + red light – to stimulate AWC only, AIA only, and AWC and AIA together, respectively (Figure 1D). AIA has variable responses to odor ^18^, so the odor-only phase was sometimes redundant with odor + red light when AIA was responsive to odor.

Altogether, these three sources of inputs (AWC, AIA, and motor activity) are integrated by AIB, and are often represented in AIB’s activity (Figure 1D). AIB is primarily influenced by motor inputs from AVA, which corroborates previous findings ^9,11,12^, with sensory inputs having a mild effect on AIB activity.

### AIB dynamics can be modeled as a convolution of AVA activity

Due to the dynamic nature of AIB’s activity in response presynaptic input, we sought to find an appropriate method to model the contributions of each neuron to AIB’s activity. The convolution of kernels with neuronal inputs is a common method to model the response properties of neurons, either as a convolution of sensory inputs ^26^ or motor behavior ^12^. The kernel itself represents the response kinetics of the neuron. Since AIB activity is tightly coupled to reversal motor activity ^9–12^, we first modeled AIB as a convolution of motor activity (Figure 2):

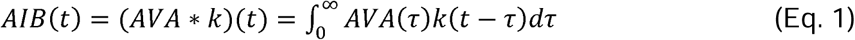

**Figure 2:**
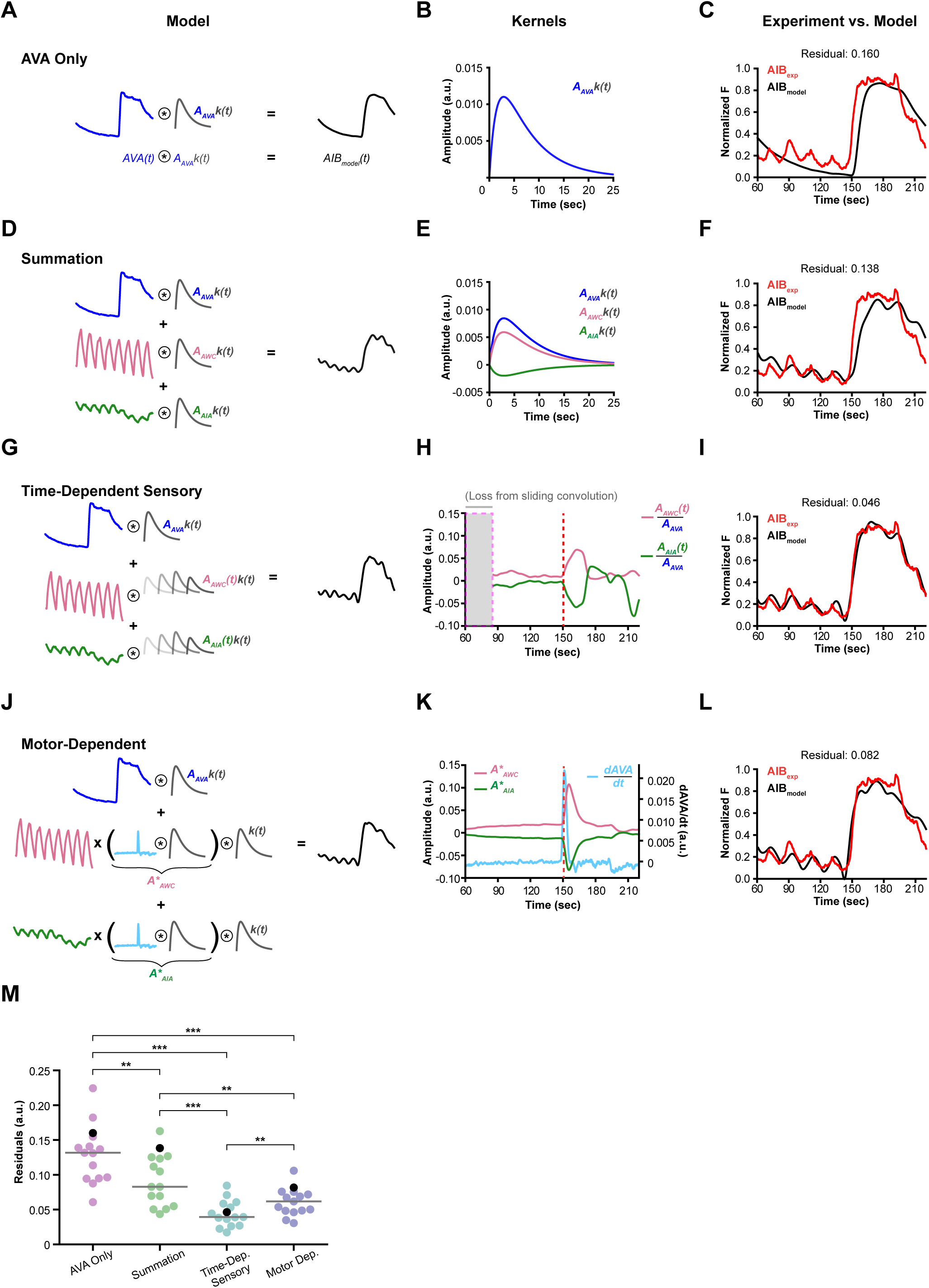
Sensory gain increases during reversal initiation. Comparison of different convolution modeling strategies. All models used odor + red light experiments for maximal AWC and AIA activation, and all example traces are from the same dataset from Figure 1D for continuity and comparison. **A)** Schematic of previous AVA-only modeling strategy, **B)** an example kernel (from Figure 2F), and **C)** resulting model with its residual (from Figure 2D, simplified for AIB and AIB model comparison). **D – F)** AVA + AWC + AIA summation model, kernels, and resulting model. **G-I)** Time-dependent sensory kernel model. **G)** Modeling strategy where the sensory kernels are time-dependent. **H)** AWC and AIA time-dependent kernel amplitudes normalized by AVA’s kernel amplitude. Sliding the convolution over time results in a loss in the beginning of the trace from the sliding window duration (indicated by shaded gray area). The dotted red line denotes onset of AVA activity in the trace. **I)** Resulting model comparison. **J-L)** Motor-dependent sensory kernel model. **J)** Sensory kernel amplitudes are weighted by 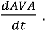 **K)** AWC and AIA kernel amplitudes are convolutions of 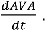 **L)** Resulting model comparison. **M)** Quantification of model performances (n=15 worms). Black dots highlight the specific traces shown throughout the figure (**C**, **F**, **I**, and **L**). P-values calculated with the Wilcoxon signed-rank test and corrected with Holm-Bonferroni; >0.05: N.S., <0.05: *, <0.01: **, <0.001: ***.

where the kernel, *k*, is:

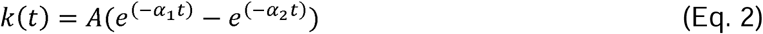

The kernel characterizes the response kinetics of AIB, where the difference between the two exponential functions defines the on-off kinetics of AIB’s response to input.

Kernel parameters were fit individually for each experiment. Convolving with AVA alone captured most AIB dynamics (Figure 1A-C, Supplementary Figure 1), while convolving with AWC or AIA alone captured very little (Supplementary Figure 2). The distributions of alpha values for the AVA convolution were fairly narrow, while the amplitudes had greater variance, consistent with response kinetics of AIB being fairly robust across experiments and animals, but the relative gain of AVA varying across trials. (Supplementary Figure 1). Due to the narrow variance of the alpha values, we chose to fix *α_1_* and *α_2_* to their median values for subsequent modeling, but allowed the kernel amplitudes to float to reflect the variable gain of AVA across animals and trials.

### AIB integration of sensory inputs is dependent on motor activity

The simplest approach to include sensory input in the model was to model AIB as a sum of convolutions (Figure 2D):

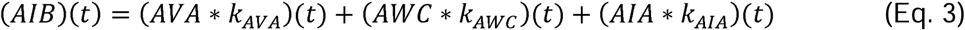

where each kernel (*k*) had the same kinetic properties (i.e. same *α_1_* and *α_2_*), but different amplitudes to reflect their differing contributions to AIB:

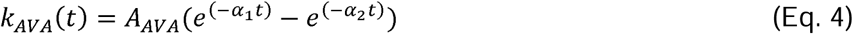

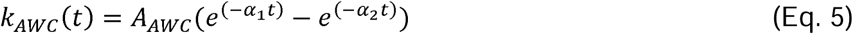

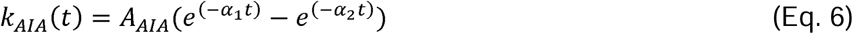

The resulting model can be rewritten as:

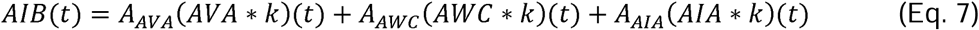

In most cases, AVA had the largest amplitude, reflecting the observation that AVA has the greatest influence on AIB activity (Figure 2E, Supplementary Figure 3). Following AVA, AWC usually had the second highest contribution, reflecting the frequent sensory dynamics provided by AWC. The contribution from AIA was typically small and negative, which is consistent with AIA having a positive response to odor which is anti-correlated with AWC and AIB’s typical response ^9,18,27^. The small contribution of AIA was surprising given its numerous synapses onto AIB. When comparing the AVA-only convolution versus the AVA+AWC+AIA summation convolution, the summation model is a significant improvement and captures the nuances of AIB activity in response to the pulses of sensory stimuli (Figure 2F, 2M).

Since the kernel amplitudes are constants, the model assumes the contribution of each neuron is static. To investigate the potential time-dependency of sensory input onto AIB, the sensory kernel amplitudes were allowed to float over time, while AVA’s amplitude remained fixed (Figure 2 G-I, Supplementary Figure 4):

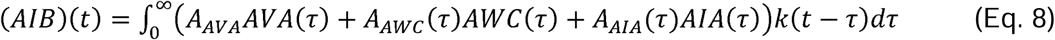

The residual of this model relative to the experimental AIB data was improved (Figure 2I,M), but also expected, since this model was over-fit for each time-point. With this approach, the amplitudes of AWC and AIA relative to AVA were small (<10%), but occasionally these amplitudes increased (Figure 2H, Supplementary Figure 4). However, we noticed that these sudden spikes in AWC and AIA kernel amplitudes frequently occurred when AVA activity increased. This seemed to indicate that the fluctuations in sensory kernel amplitudes were not arbitrary, but tied to motor activity.

Since the sensory kernel amplitudes correlated with sudden changes in motor activity (Figure 2H), we modeled the sensory kernel amplitudes as convolutions of the time-derivative of motor activity (Figure 3J):

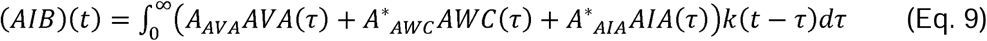

**Figure 3:**
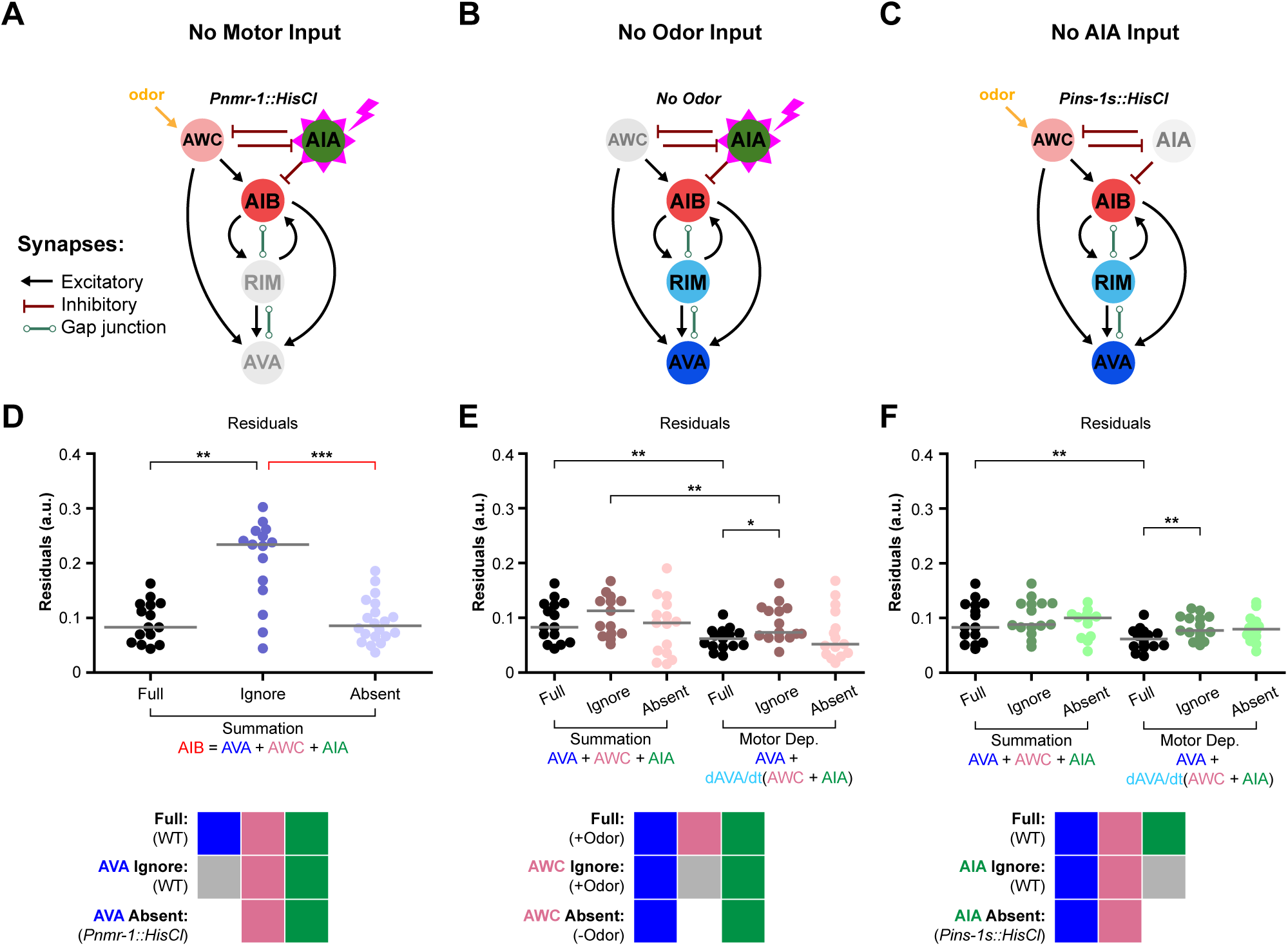
Individual contributions of sensory and motor neurons. Schematics for removing the activity of motor input **(A)**, AWC **(B)**, or AIA **(C)** in our neuronal network. **D-F)** Resulting residuals of models in different experimental contexts. “Full” models denote the presence of all three neurons (AVA, AWC, AIA) for either summation or derivative models. “Ignore” models do not include specific neurons in models for experiments where all neurons were active. “Absent” models lack neuronal activity for specific neurons. Block diagrams: Full color = neuron active and included in model, Gray = neuron active and not included in model, White = neuron silenced. P-values calculated using either Wilcoxon signed-rank test between “Full” and “Ignore” models (black bars) or Mann-Whitney U test for any comparisons between “Absent” models and other models (red bars). Significance corrected using Holm-Bonferroni; N.S.: >0.05, <0.05: *, **: <0.01, ***: <0.001. n = 15 worms for all “Full”, “Ignore”, and AWC “Absent” data; n = 20 for *Pnmr-1::HisCl* “Absent”; and n = 11 for *Pins-1s::HisCl* “Absent”.

Where:

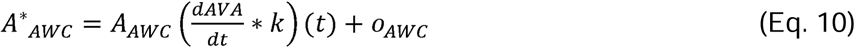

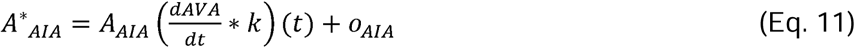

With this approach, the sensory kernel amplitudes are time dependent, but not arbitrary. Instead, they are tied to changes in motor activity (Figure 2K). The resulting model (Figure 2L) performs similarly to the overfit time-dependent sensory model, albeit slightly worse, yet still significantly better than the summation model (Figure 3M). It should be noted that despite the worsened performance of the motor-dependent model in comparison to the time-dependent sensory model, the time-dependence of the motor-dependent model is not arbitrary. The improved performance is not simply due to the structure of the model. Swapping the dependencies (i.e. tying motor gain to changes in sensory dynamics), or modeling AIB as a sum of motor and motor time-derivatives did not improve performance (Supplementary Figure 5).

Tying the sensory amplitudes to the derivative of AVA activity was sufficient to produce comparable performance to the overfit model. This implies that sensory gain increases while the worm performs reversals.

### Silencing individual neurons does not alter model performance

The prior models were based on experiments where sensory and motor neurons were active. To investigate individual contributions of each neuron, we modeled AIB activity in experiments where AWC, AIA or AVA were individually silent. For AIA or the motor neurons, a histamine-gated chloride channel (HisCl) ^28,29^ was expressed in these neurons individually to chemogenetically silence them (Figure 3A, C). For AWC, this was done by simply not providing odor stimuli (Figure 3B).

We used two different methods to compare the contributions of individual neurons, which we dubbed “ignore” versus “absent”. In our “full” models, all three presynaptic neurons were experimentally active, and all were included in the models (Figure 3D – 3F). In our “ignore” models, these same full experiments were modeled but with one neuron absent to compare how well the model performed if one neuron’s contribution was ignored. In contrast, in our “absent” models, a neuron was experimentally silenced, and the model excluded this neuron’s contribution. If a presynaptic neuron contributes significantly to AIB’s activity, then the “ignore” models should perform poorly compared to the “full” model. If a presynaptic neuron significantly alters the integration properties of AIB, then the “absent” models should perform worse than the “full” models.

Models where AWC or AIA were ignored only performed slightly worse for the motor-dependent model, but not for the summation model (Figure 3B-C,E-F). However the magnitude of the effect was not as severe as ignoring motor input (Figure 3D). This suggests that the sensory neurons do play an important role in the network but their individual contributions are slight. This is consistent with how the models performed when either AWC or AIA was silent in the context of motor activity. Removing either neuron did not significantly affect the performance of the summation or motor-dependent models (Figure 3E-F). This was largely due to how redundant the information from these neurons were. If AWC was ignored or silent, the amplitude of the AIA kernel increased to compensate for the loss of AWC (Supplementary Figure 3).

As expected, ignoring the motor contribution to AIB’s activity led to poor model performance which now solely relied on the sensory input (Figure 3D). However, if motor activity was silenced in the “absent” model, this sensory-only model performed as well as the “full” model, confirming that in the absence of motor activity, AIB activity can be modeled as a sensory-only interneuron. However, it should be noted that the relative amplitude of AIB activity was small compared to the fluorescence changes observed when motor activity was present (Supplementary Figure 6), which is consistent with the relatively small contribution sensory input makes to AIB activity. Also, the relative amplitudes of the AWC and AIA kernels were comparable to the baseline amplitudes of the motor-dependent model (Supplementary Figure 3), which is expected if there is no motor contribution. In the absence of motor input, the motor-dependent model essentially becomes a summation model. However, the relative contribution of sensory inputs to AIB remained low.

Overall, these silencing experiments provide an orthogonal approach to confirm the contributions of sensory and motor inputs to AIB activity. Ignoring motor input dramatically decreases model performance. However, removing motor input does not necessarily lead to worse model performance since AIB activity is now largely tied to sensory input. AIB activity decreases due to the lack of motor dynamics, which negates the motor-dependence in sensory gain. Sensory information from AWC and AIA is largely redundant, and ignoring one neuron can be compensated by increasing the gain in the other neuron.

### Motor-dependent model outperforms summation model for whole-brain NEUROPAL data

While the activity of sensory and motor neurons explored here are sufficient to explain most AIB activity, there are still numerous other AIB presynaptic partners. To explore AIB integration in the context of the whole brain, we turned to whole-brain *C. elegans* recordings with consistent sensory stimulation ^30^. This dataset encompasses the calcium activity of 231 neurons (189 in the head, 42 in the tail) in 21 worms in response to chemosensory stimuli. In these recordings, ten-second pulses of three chemicals were delivered in a randomized order: 2,3-pentanedione (an attractive odor), 2-butanone (an attractive odor), and sodium chloride (a repulsive taste stimulus). This dataset was chosen over other whole-brain worm datasets ^12,31^ for two reasons. One, this particular dataset had consistent pulses of sensory stimulation, similar to our own dataset, so sensory neuron dynamics were well-represented. Two, AIA is often under-represented in these datasets due to weak calcium dynamics in the soma, but this dataset had several AIA recordings. A different GCaMP was used (GCaMP6s), but the kinetic differences between GCaMP6s and GCaMP7s are small compared to the response dynamics of AIB^24,32^. Due to the sensory-driven nature of this dataset, we investigated whether AIB exhibited the same motor-dependent integration for other sensory neurons other than AWC.

We first compared the summation and motor-dependent models for AWC, AIA, AVA neuronal activities from the NeuroPAL dataset. We initially convolved AVA with our kernel to model AIB activity (Figure 4A). We found the kinetic parameters for the AIB kernel from the NeuroPAL data were similar to our own (our median values α_1_: 0.16 sec^-1^, α_2_: 0.64 sec^-1^; NeuroPAL α_1_: 0.15 sec^-1^, α_2_: 0.72 sec^-1^), which suggests the observed response kinetics from both experiments is an innate property of AIB (Supplementary Figure 7). As observed in our own experiments, AIB activity is largely driven by motor input (Supplementary Figure 8). We next applied the same summation model, this time incorporating both AWC OFF and AWC ON activity from the NeuroPAL dataset (Fig 4A, Supplementary Figure 8). In our experiments, we only recorded AWC OFF responses to isoamyl alcohol using the *str-2* promoter, but both AWC OFF and AWC ON respond to IAA and therefore should both contribute to AIB activity ^33,34^.

**Figure 4:**
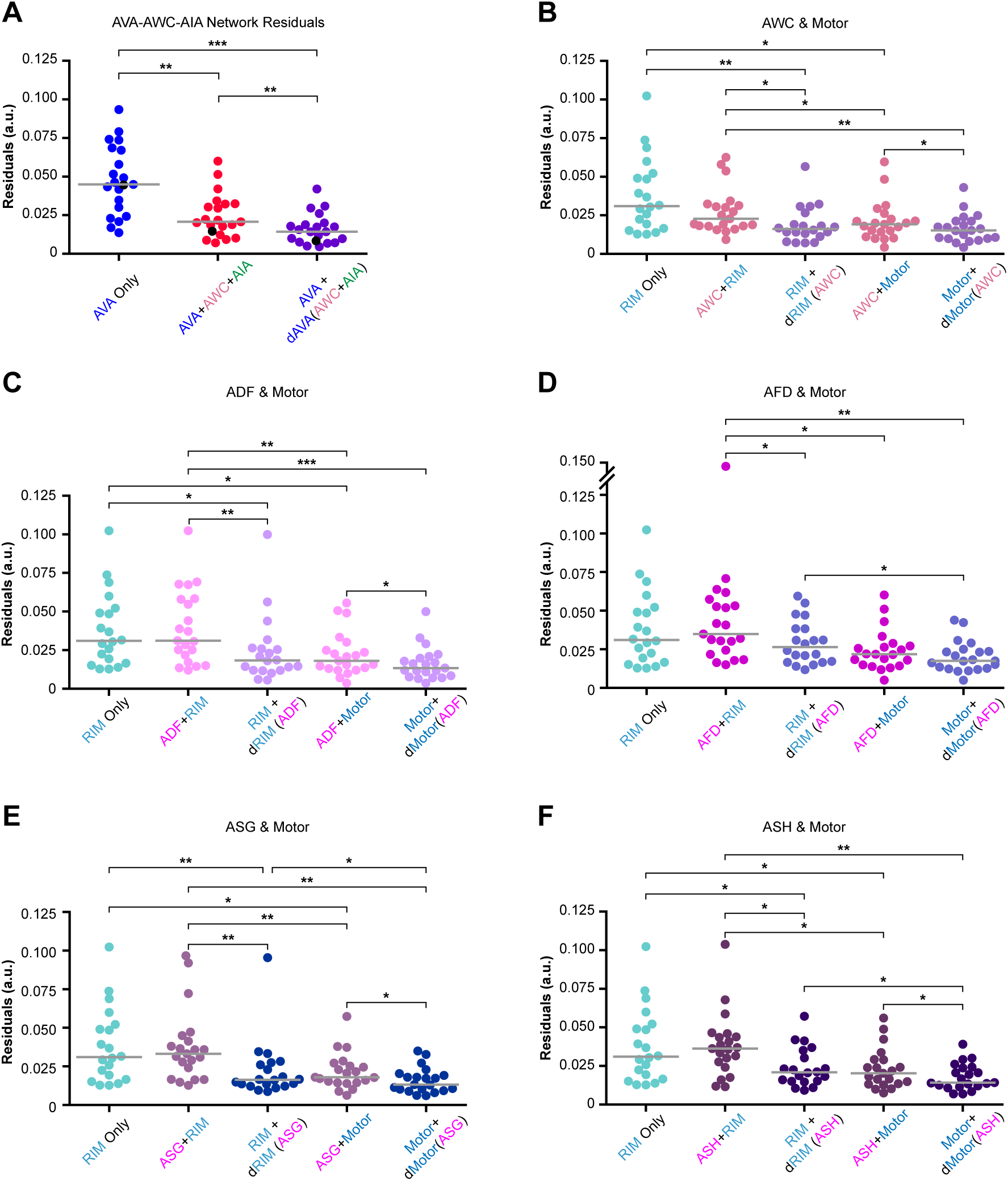
Sensorimotor integration for sensory neurons from whole brain recordings. **A)** Residuals for AIB models for AVA-only, sum of AVA, AWC, and AIA, or AVA-dependent sensory gain. **B-F)** Residuals of AIB models for RIM-only, sensory neuron + RIM, RIM-dependent sensory gain, sensory neuron + motor inputs, and motor-dependent sensory gain. **B**) AWC (using individual AWC OFF and AWC ON traces), **C**) ADF (using individual L-R traces), **D**) AFD (using averaged L-R trace), **E**) ASG (using individual L-R traces), or **F**) ASH (using averaged L-R trace). n = 20 worms for RIM-only residuals, as only 20 recordings are available, and n = 21 worms for sensory-motor residuals. P-values calculated with the Wilcoxon signed-rank test and corrected with Holm-Bonferroni; >0.05: N.S., <0.05: *, <0.01: **, <0.001: ***.

Although the NeuroPAL experiments did not use IAA, AWC OFF responds to 2,3-pentanedione, and AWC ON responds to 2-butanone ^33,34^, which were both delivered during imaging, mirroring a similar pattern of AWC stimulation in our own experiments. As observed in our experiments, incorporating sensory information improved model performance, consistent with AIB’s role in integrating sensory information (Figure 4A). Additionally, consistent with our observations, the motor-dependent model outperformed the summation model for the NeuroPAL data (Fig 4A, Supplementary Figure 8).

We then wanted to extend our modeling approach by incorporating more sensory inputs to investigate how AIB integrates information from the whole brain. We selected the most relevant presynaptic sensory inputs based on general correlation to AIB activity – AWC, ADF, ASG, ASH, AFD. Sensory neuron activities were highly variable since they responded differently to the three chemicals used in the Yemini *et. al.* experiments. To assess how motor activity may influence the gain of these sensory inputs for AIB, we tested each neuron individually as a sum of RIM activity and the sensory neuron’s activity, or as a function of the derivative of motor (RIM) activity, as used previously for AWC. Consistent with our prior observations with AWC, models that tied the gain of sensory neuron activity to changes in motor activity improved model performance (Figure 4B-F). This was true even for sensory neurons that responded to different stimuli, such as ASH and AWC.

Other motor and motor interneurons in addition to RIM, principally SAAV and RIB, also synapse on to AIB. To include their contributions, a “motor” signal was constructed as the sum of their activities (thought inverted for RIB since it anti-correlates with AIB). This motor signal, and its derivative, was used in place of RIM for the summation and motor-dependent models. Including these neurons improved model performance, likely due to the additional information these neurons provide. RIB activity is tied to forward motor responses, and SAAV are head motor neurons. For most sensory neurons tested, the motor-dependent model out performed the summation model, consistent with observations made for single motor neurons. AIB activity is primarily influenced by motor activity, and while sensory input also contributes, its gain is tied to changes in motor activity.

## Discussion

AIB is an important interneuron that integrates both motor and sensory information, and is critical to behavioral responses to sensory stimuli. AIB drives reversal behavior, but is also activated by reversal interneurons, creating a positive feedback loop. While much of its activity is dominated by the reversal motor-circuit, sensory triggered responses can also be observed, however these responses are more variable, and appear to be tied to the state of the motor-circuit ^9,10^. The nexus of sensory and motor input onto AIB, and its role in reversal behavior, make it an ideal candidate for sensory-reinforcement. The removal of attractive stimuli increases the probability of reversals as the worm tries to reorient and find the attractive cue. AWC is stimulated by the removal of odor, and has excitatory synapses onto AIB, while AIA is stimulated by attractive odor, and has inhibitory synapses onto AIB^8^. This is consistent with driving reversal behavior when the environment decreases in quality. Since AIB is also activated by the reversal circuit which it contributes to, it can serve as a positive reinforcer of reversal behavior when it matches sensory information. This state-dependent sensory response, along with the biased synaptic anatomy of the AIB neurons, led us to investigate how this neuron explicitly integrates presynaptic input.

Sensory input influences AIB activity, though AWC and AIA input appeared to be redundant, and silencing AIA did not have a strong impact on AIB activity. The role of AIA in the circuit may be more context-dependent over observation periods not measured here. Prior work has shown that AIA can operate on different timescales, and can promote roaming or dwelling states depending on sensory context ^27^.

While AIB acivity is largely driven by the reversal motor circuit, we found the onset of reversal behavior increased the gain in sensory input. This was observed in our experiments, as well as for other presynaptic sensory neurons in other experimental datasets under different contexts. AIB does integrate sensory information, but the gain in this information increases when the worm initiates a reversal. This is a useful feature for behavioral reinforcement when the worm alters its behavior. For AIB, the saliency of sensory input is highest during when a reversal is initiated, and helps reinforce the decision to reverse. Sensory-motor feedback between motor states and thermotaxis has been previously observed as well ^35^, suggesting this approach can be extended to other sensory modalities as well. Motor-dependent sensory responses have also been observed in the touch response, where mechanosensory responses are dependent on the locomotion state of the animal ^22,23^. Associating changes in sensory perception to changes in motor activity is also a critical feature of corollary discharge: the need to cancel sensory input generated by self-motion ^36^. It is advantageous to consider context when assessing the saliency of sensory input.

Context-dependent sensory responses are common in nervous systems besides the worm. In the mouse visual cortex, visual gain is enhanced during locomotion compared to remaining stationary ^37–39^. In *Drosophila* larvae, animals execute a different series of runs, turns, and head sweeps during positive or negative thermotaxis ^40^. Corollary discharge is observed in most sensorimotor systems, as the need to differentiate changes in sensory perception due to egocentric motion and the environment is critical to navigation ^41^. Ethologically, context-dependent sensory responses are important for evaluating the value of the environment. Animals constantly assess internal cues with sensory information to inform their behavioral decisions.

Anatomically, the separate computations performed by AIB are reflected in its anatomy. While the presynaptic inputs along its process are integrated, the sensory input on the proximal process is conditionally integrated based on motor activity from the distal process. The cellular basis of this remains to be determined. Other interneurons such as RIA not only have segregated presynaptic inputs, but also produce compartmentalized calcium dynamics along its process, despite an absence of a membrane barrier between these compartments ^42^. This property has also been observed in the RIS neuron ^43^. Other neurons besides these have clustered synapses along their processes, and may also be sources of nonlinear computation ^5^.

Computationally, how does a neuron compute this context-dependence? Here we model context-dependence of sensory gain as a non-linear function that is simply the product of sensory input with the derivative of the motor response. In other words, the gain in sensory input increases when there is a change in motor activity. While neuron polarization is an additive result of presynaptic excitation and inhibition, non-linear summation has also been observed.

Two mammalian examples are superlinear and sublinear visual-auditory integration in the mouse superior colliculus ^44^, and orientation selectivity of layer 2/3 neurons in the visual cortex of ferrets ^45^. In these two systems, the inclusion of nonlinear summation allows for greater tuning to their respective stimuli. Our results indicate nonlinear computation may also occur in the *C. elegans* brain. This observation aligns with previous reports stating that global switching between behavioral states is accomplished through nonlinear rather than linear dynamics (Morrison et al. 2021). Whole-brain imaging in the worm has shown a considerable degree of state-switching that is tied to behavioral dynamics that seem to show signatures of deterministic chaos ^11^. Such a system cannot emerge from a solely linear system; amplifying non-linearities are necessary to destabilize the system. The non-linear gain modeled here would be an example of just such a nonlinearity. If such computations are coupled to feedback in the system, they could be a source of deterministic chaos, and drive the highly variable behavioral switching observed in *C. elegans* ^11^.

## Materials and Methods

### Worm Maintenance and Preparation for Imaging

Worms were grown and maintained under standard conditions using 6 cm nematode growth medium (NGM) plates and fed *Escherichia coli* OP50 bacteria ^47^. Worms were picked at the L4 stage and kept at 20°C 16-24 hours before all imaging sessions so that they would be young adults during imaging. L4 worms were placed onto either OP50 lawns, or onto retinal plates if the worm expressed Chrimson ^25^. To make retinal plates, all-trans retinal (Sigma-Aldrich) was mixed with OP50 (50 μM) and was seeded a few hours in advance before picking L4 worms expressing Chrimson. All retinal plate handling was performed in the dark to minimize photooxidation.

### Worm Strains

AGG0118 and AGG0041 strains were generated by injecting respective constructs below into wild-type N2 worms using standard microinjection protocol ^48^. CX17432 worms were obtained from ^18^, and expressed two co-injection markers – *Pelt-2::mCherry* (intestine) and *Punc-122::dsRed* (coelomocyte) – to indicate expression of AIA::GCaMP5A and AIA::Chrimson, respectively. To generate AGG0116, we selected CX17432 worms that only expressed the intestinal marker to allow us to select for AIA::Chrimson expression to cross into our AVA, AIB, AIA::GCaMP7s line (AGG0118) which already had GCaMP7s expression in AIA. AGG0142 was generated by injecting the HisCl construct and the co-injection marker *Punc-122::dsRed* into AGG0118, and AGG0145 was similarly generated by injecting into AGG0116. We used different combinations of two AIA promoters, *Pins-1(s)* and *Pgcy-28d*, in our strains. *Pins-1(s)* is more specific to AIA but less bright, which was useful for histamine silencing and targeting optogenetically with Chrimson. *Pgcy-28d* has more non-specific labeling of other neurons other than AIA but has brighter expression, which was useful for calcium imaging with GCaMP7s.

| Strain ID | Source | Expression | Genotype/Construct Information | Figure Appearances |
| --- | --- | --- | --- | --- |
| AGG0041 | This paper | GCaMP7s in AWC. Co-injection expression: red coelomocytes. | *Pstr-2::NLS::GCaMP7s* = 30 ng/ul, *Punc-122::dsRed* = 15 ng/ul | Figures 1-4 |
| AGG0118 | This paper | GCaMP7s in AVA, AIB, and AIA. Co-injection expression: green coelomocytes. | *Prig-3::NLS::GCaMP7s* = 40 ng/ul, *Pinx-1::GCaMP7s* = 50 ng/ul, *Pgcy-28d::GCaMP7s* = 50 ng/ul, *Punc-122::GFP* = 10 ng/ul | Figures 1-4 |
| CX17432 | Obtained externally <sup>18</sup> | GCaMP5A and Chrimson in AIA. Co-injection expression: red coelomocytes and red intestines. | *Pgcy-28d::GCaMP5A* = 50 ng/ul, *unc-122::dsRed* = 15 ng/ul, *Pins-1(s)p::Chrimson::sl2::mCherry* = 20 ng/ul, *Pelt-2p::mCherry* = 2 ng/ul | Figures 1-4 |
| AGG0116 | This paper | GCaMP7s in AVA, AIB, and AIA, and Chrimson in AIA. Cross between AGG0118 and CX17432. Co-injection expression: green coelomocytes and red intestines. | *Prig-3::NLS::GCaMP7s* = 40 ng/ul, *Pinx-1::GCaMP7s* = 50 ng/ul, *Pgcy-28d::GCaMP7s* = 50 ng/ul, *Punc-122::GFP* = 10 ng/ul, *Pins-1(s)::Chrimson::sl2::mCherry* = 20 ng/ul, *elt-2p::mCherry* = 2 ng/ul | Figures 1-4 |
| AGG0142 | This paper | Histamine-gated chloride channel (HisCl) expression in AIA, and GCaMP7s in AVA, AIB, | *Pins-1(s)::HisCl::mCherry* = 100ng/uL, *Prig-3::NLS::GCaMP7s* = 40ng/ul, *Pinx-1::GCaMP7s* = 50ng/ul, *Pgcy-28d::GCaMP7s* = | Figure 4 |
|  |  | and AIA. Co-injection expression: red and green coelomocytes. | 50ng/ul, *Punc-122::dsRed* = 10ng/uL, *Punc-122::GFP* = 10ng/ul |  |
| AGG0145 | This paper | HisCl expressed in motor neurons such as RIM and AVA, Chrimson expressed in AIA, and GCaMP7s in AVA, AIB, and AIA. | *Pnmr-1::HisCl::mCherry* = 50ng/uL, *Prig-3::NLS::GCaMP7s* = 40ng/ul, *Pinx-1::GCaMP7s* = 50ng/ul, *Pgcy-28d::GCaMP7s* = 50ng/ul, *Punc-122::GFP* = 10ng/ul, *Pins-1(s)::Chrimson::sl2::mCherry* = 20ng/ul, *Pelt-2p::mCherry* = 2ng/ul | Figure 4 |

### Live Calcium Imaging using Microfluidic Chips

GCaMP animals were imaged using microfluidic chips ^20^ to monitor calcium activity. Worms were restrained in the chip during imaging and were also paralyzed with 10 mM tetramisole hydrochloride (Sigma-Aldrich) dissolved in S-basal buffer to minimize movement during imaging. Young adult animals were submerged in a tetramisole bath 10-15 minutes before imaging to ensure paralysis. Each animal was imaged for the duration of all 3 phases (odor, red light, and odor + red light; 2300 frames per phase) of an experiment, up to 2 times. Odor provided was 100 μM isoamyl alcohol, and odor delivery was controlled using LEE valves and AutoMate Scientific Valve Link 8.2. Each phase and set of experiments were imaged within a minute of each other to readjust the focal plane as needed. To record from neurons, we focused on the cell body of AWC in strain AGG0041, and in any strain expressing GCaMP in AVA-AIB-AIA we focused on the cell bodies of AVA and AIB and the process of AIA all in the same focal plane. We focused on AIA’s process as it has higher calcium dynamics compared to its soma.

### Microscopy with Light Strobing

Microscopy was performed using a Nikon Eclipse Ti2 inverted microscope. Videos were recorded at 10 frames per second using a 40x water objective (Nikon N Apo LWD 40x / 1.15 WI λS). Strobing was performed using Nikon NIS Elements software using a custom Illumination Sequence protocol to minimize photobleaching and oversaturation of GCaMP7s signals, at a rate of 5 ms of blue light (470 nm, offset by 2 ms beforehand) per 100 ms cycles.

### Optogenetic Stimulation

To stimulate AIA::Chrimson, we used an external red LED (Mightex 617 nm). We employed a strobing approach to minimize the activation of Chrimson during GCaMP imaging due to spectral overlap between GCaMP and Chrimson. Acquisition was at 10 Hz with the following illumination protocol: 5 ms blue light to excite GCaMP, 1 ms dark, 93 ms red light to activate Chrimson, 1 ms dark. For periods when Chrimson was not activated, illumination was 5 ms of blue light, followed by 95 ms of dark. This allowed for any blue-light photoactivation of Chrimson to decay during non-stimulation periods.

### Histamine-Gated Chloride Channel Silencing and Imaging

A histamine-gated chloride channel (HisCl) was expressed in either AIA under the *Pins-1(s)* promoter, or in several command neurons (RIM, AVA, AVB, AVD, AVE, PVC, AVG, AVF) ^49,50^ under the *Pnmr-1* promoter. Neurons were silenced with the addition of 10 mM histamine dihydrochloride (Sigma-Aldrich) in loading buffer and all liquids used during microfluidic chip imaging. Worms were silenced for 20 minutes prior to imaging.

### Data Processing

We individually analyzed each neuron using custom ImageJ and Python scripts. Regions of interest were hand-annotated to quantify neuron fluorescence and background. Neuron traces were first background subtracted, and then the first 350 frames (35 seconds) of each phase per video were cut off to remove initial photodecay artifacts (Fig 1D). Then, we normalized all 3 phase recordings for each worm across a set of experiments. We did this to ensure that we captured the full dynamic range per set of experiments for each neuron, since some phases were designed to not activate specific neurons. Normalization was performed by taking the F_max_ and F_min_ of each set of experiments per worm, not for individual phases, and then calculating 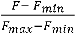 so that each neuron trace is normalized from 0 to 1. These normalized traces were used for all convolution modeling experiments.

### Modeling

Convolution modeling was done using custom Python scripts. To find the model of best fit for each analysis we used the SciPy minimize function with the Nelder-Mead method, which utilizes a simplex approach. We calculate the difference between modeled AIB and real AIB using a cost function, such that a smaller residual denotes a better fit:

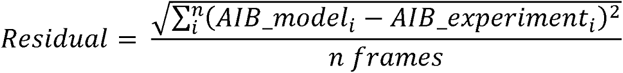

### Modeling – AVA-Only

Kernel parameters were optimized by gradient descent. Initially the parameters were optimized separately for each kernel, but since the *α_1_, α_2_* parameters were stable across experiments, all models used the median *α_1_, α_2_* values. Amplitudes (*A*) were fit separately for each recording.

### Modeling – Summation

The amplitudes for all kernels were optimized individually for each recording. The same kinetic parameters (*α_1_, α_2_*) from the AVA-only modeling were used for all modeling.

### Modeling – Time-Dependent Sensory Inputs

For the sliding window used in Figure 3G – 3I, a sliding window of convolutions over 50 second intervals was passed over the entire trace. During this short convolution window, the amplitudes of both AWC and AIA’s kernel were fitted (*A_neuron_*(*t*)). Note that A_AVA_ was a fixed value; only A_AWC_ and A_AIA_ were allowed to float over time. This process generated a series of short, convolved model fragments of the entire trace, which was then averaged to provide the models in Figure 3 G-I, M.

### Modeling – Motor-Dependent

The amplitudes of AWC and AIA were a function of a convolved derivative of the AVA activity. An offset term (*o*) was added to account for sensory input in the absence of motor activity.

### Modeling with NeuroPAL data

Data provided courtesy of the Yemini lab ^28^. For the sake of comparative analysis and consistency, we applied the same metrics of modeling our data to the NeuroPAL dataset. This included normalizing traces from 0 to 1, cutting off the beginning of the traces to exclude initial imaging artifacts, using a 25 second (100 frame) kernel length (4 fps rate for the head neurons), and using fixed median α_1_ and α_2_ values based on the AVA-only model.

### Modeling with NeuroPAL data – AVA-AWC-AIA Network

We combined AVA L/R and AIA L/R into their own averaged traces due to similarity in activity, and AWC ON/OFF were used separately.

### Selecting and consolidating relevant AIB inputs in NeuroPAL

We first documented all of AIB’s presynaptic inputs and gap junction partners present in the NeuroPAL dataset based on ^5^ which ended up totaling 39 inputs to AIBL and 37 inputs to AIBR. Since many of the presynaptic inputs had polymodal functions (i.e. RIM is a motor interneuron), we categorized each input based on what non-interneuron function it had (i.e. RIM was considered a motor input). If the neuron was solely an interneuron, it was categorized as such. Because not all presynaptic inputs necessarily influence AIB activity, and trying to apply our modeling methods to a large dataset risks overfitting parameters, we first narrowed down this list to what would be the most relevant inputs of AIB activity. We compared the activity of each presynaptic neuron and ranked how well they correlated to AIB L-R activity using Pearson’s correlation (ρ). We then applied an arbitrary cutoff of ±0.3 to rule out the least correlative neurons. There were also some trials out of the total of 21 worms where some neurons had very few recordings, so those with less than 10 recordings were excluded as well. We then further ruled out neurons that only had inputs onto one of the AIB pairs but not both left and right. Finally, many of the presynaptic inputs had left-right pair symmetry in activity that could further be consolidated into one representative activity by taking the mean of the left and right pairs. We decided which neuron left-right pairs could be consolidated into one based on their ρ value, where a value above 0.75 would be combined. AIB L-R itself was also combined as an averaged trace for modeling. This full process of consolidation allowed us to narrow our list from 39 (AIBL) and 37 (AIBR) neurons to 14 of potentially the most relevant presynaptic inputs influencing AIB activity.

### Modeling with NeuroPAL data – Individual sensory neurons and motor activity

We considered presynaptic inputs to AIB present in the NeuroPAL dataset, and based on correlated activity to AIB, we found 5 sensory neurons (ADF, AFD, ASH, ASG, and AWC) and 3 motor-related neurons (RIM, SAAV, and RIB). We used these for summation and derivative modeling using individual sensory neurons with only RIM or with the collective summed activity of the motor neurons. RIM and SAAV are associated with reversal behavior, and due to RIB’s association with forward movement, we used inverted RIB activity. Since RIM synapses onto AIB, we used RIM-only activity instead of AVA. We used the resulting RIM/motor amplitudes from the summation models as fixed parameters in the corresponding derivative models. This was to be consistent with our own data where we used the same AVA amplitude from the AVA+AWC+AIA models and fixed it in the derivative model. For modeling with the motor derivative, we used the average of the derivatives of all motor neurons, as they exhibited similar activity and derivatives.

---

**<u>Supplementary Figure 1: A convolution of AVA activity alone is sufficient to capture most AIB dynamics.</u>**

**A – C)** AVA convolution model of AIB for each phase from a single worm experiment. **D – F)** Resulting kernels from the models shown in A – C. **G – I)** Resulting kernel parameters from all all traces (n= 15 worms), where each kernel was uniquely fit for **D)** amplitude, **E)** alpha 1 (*α_1_*), and **F)** alpha 2 (*α_2_*). Medians represented by black stars.

**<u>Supplementary Figure 2: Single-neuron input modeling of AIB activity</u>. A)** Residuals of single-neuron input models for AIB during the three experimental contexts (n = 15 worms). **B – D)** Example traces of AIB, AVA, and AWC for AWC-only models of AIB activity. **E – G)** Example traces of AIB, AVA, and AIA for AIA-only models of AIB activity. P-values calculated with the Wilcoxon signed-rank test and corrected with Holm-Bonferroni; >0.05: N.S., <0.05: *, <0.01: **, <0.001: ***.

**<u>Supplementary Figure 3: Kernel amplitudes in different model contexts</u>. A)** AVA kernel amplitudes for summation models that included AVA, AWC, AIA (Full), ignored AIA (iAIA), silenced AIA (-AIA), ignored AWC (iAWC) or silenced AWC (-AWC). **B)** AWC kernel amplitudes for summation models that included AVA, AWC, AIA (Full), ignored AVA (iAVA), silenced AVA (-AVA), ignored AIA (iAIA) or silenced AIA (-AIA). **C)** AIA kernel amplitudes for summation models that included AVA, AWC, AIA (Full), ignored AVA (iAVA), silenced AVA (-AVA), ignored AWC (iAWC) or silenced AWC (-AWC). **D)** AWC and AIA kernel amplitudes for motor-dependent models where AWC, AIA, and AVA are convolved to model AIB activity (Full), only AVA and AWC (iAIA), only AVA and AIA (iAWC), AIA is silenced (-AIA), or AWC is silenced (-AWC). **E)** AWC and AIA kernel offsets for motor-dependent models where AWC, AIA, and AVA are convolved to model AIB activity (Full), only AVA and AWC (iAIA), only AVA and AIA (iAWC), AIA is silenced (-AIA), or AWC is silenced (-AWC). **F)** Comparison of kernel offsets from motor-dependent model (*O_neuron_*) with sensory neuron amplitudes from motor-silenced experiments (*A_neuron_, -AVA*). n = 15 worms for all “Full”, all ignored (iAIA and iAWC), and AWC silenced data, n=20 for AVA-silenced, and n = 11 for AIA silenced data. P-values calculated with the Wilcoxon signed-rank test (black) or Mann-Whitney U (red) and corrected with Holm-Bonferroni; >0.05: N.S., <0.05: *, <0.01: **, <0.001: ***.

**<u>Supplementary Figure 4: Sensory time-dependent model</u>. A)** The AVA kernel has a static amplitude that is passed over the entire observation window (blue). AIA and AWC each are assigned a kernel amplitude based on a short convolution window (pink, 50s). All kernels have a length of 25s (cyan). **B)** (Top row) The AWC and AIA kernel amplitudes are optimized within the sliding window, which is then shifted by one frame (100 ms), and optimized again. (Bottom row) The predicted AIB trace (black) for the short convolution window is saved. **C)** The changing kernel amplitudes for AWC and AIA are saved. **D)** The AWC and AIA kernel amplitudes for each window plotted are normalized by the AVA amplitude for comparison. **E)** The short overlapping AIB traces are averaged across the observation window. **F)** Example of overlapping AIB models from short convolutional chunks (multi-colored traces), and the average of these overlapping predictions (black). Dashed red line in **D** and **F** indicates when AVA activity rose.

**<u>Supplementary Figure 5: Alternative models where motor kernel amplitudes are</u> <u>dependent on sensory time-derivatives</u>. A)** AIB modeled as a sum of a convolution of AVA and AVA conditioned on the time-derivative of AVA (**A**-**C**). **B)** The AVA kernel plotted along the time-derivative of AVA. **C)** Predicted AIB (black) plotted with experimentally observed AIB (red). **D)** AIB modeled as a sum of a convolution of AWC and AVA conditioned on the time-derivative of AWC (**D**-**F**). **E)** The AVA kernel plotted along the time-derivative of AWC. **F)** Predicted AIB (black) plotted with experimentally observed AIB (red). **G)** AIB modeled as a sum of a convolution of AVA and AVA conditioned on the time-derivative of AIA. **H)** The AVA kernel plotted along the time-derivative of AIA. **I)** Predicted AIB (black) plotted with experimentally observed AIB (red) (**G**-**I**). **J)** Residuals from different convolutional models (n = 15 worms). P-values calculated with the Wilcoxon signed-rank test and corrected with Holm-Bonferroni; >0.05: N.S., <0.05: *, <0.01: **, <0.001: ***.

**<u>Supplementary Figure 6: Median fluorescent values for AIB</u>.** Peak (median of top 10%) fluorescent values for AIB GCaMP change in fluorescence (ΔF) across three experimental phases for WT (black) and the motor silenced background (blue, *Pnmr-1::HisCl*). n = 15 for WT worms, and n = 20 for *Pnmr-1::HisCl* worms. P-values calculated with the Mann-Whitney U test and corrected with Holm-Bonferroni; >0.05: N.S., <0.05: *, <0.01: **, <0.001: ***.

**<u>Supplementary Figure 7: Motor-dependent kernel parameters in different experimental</u> <u>contexts</u>.** AVA kernel parameters for **A)** amplitude, **B)** alpha 1 (*α_1_*), and **C)** alpha 2 (*α_2_*) for data modeled in NeuroPAL worms (n = 21 worms). Black stars are the median of each corresponding parameter.

**<u>Supplementary Figure 8: Modeling AIB integration in the NeuroPAL dataset</u>. A**) Example trace of AIB, AWC OFF, AWC ON, AIA, and AVA activity in a NeuroPAL worm. To keep analysis consistent with our own data, traces were normalized and the beginning of the trace (50 frames) was cut off due to initial photodecay. For the sake of consistency, the same trace will be shown in the rest of the figure to showcase resulting kernels and models. **B**) Schematic of AVA-only modeling, **C**) resulting kernel, and **D**) resulting AIB model versus real AIB. **E – G**) Schematic of summation model, resulting kernels, and AIB model. **H – J**) Schematic of derivative model, resulting kernels, and AIB model. **K**) Resulting residuals from each model, black dot highlights the specific model residuals shown in **D**, **G** and **J** (n = 21 worms). P-values calculated with the Wilcoxon signed-rank test and corrected with Holm-Bonferroni; >0.05: N.S., <0.05: *, <0.01: **, <0.001: ***.

## Acknowledgements

We would like to thank A. Corver for initial help with analysis coding, M. Echterling for clustering analysis to identify AIB, E. Pyfrom and J. Reategui for the generation of AGG0041, and other members of the Gordus lab for their generous support and feedback. We thank C. Bargmann for the CX17432 strain. We also thank S. Flavell for helpful comments. A.G. acknowledges funding from NIH (R35GM124883).

## Notes

### Competing Interest Statement

The authors have declared no competing interest.

