## Supplementary Table 1 for "Nonlinear integration of sensory and motor inputs by a single neuron in *C. elegans*"

| Quantity and types of synapses Along AIBL Process | | | | | | |
| --- | --- | --- | --- | --- | --- | --- |
| AIBL | **Proximal** | | | **Distal** | | |
|  | **Gap** | **Inputs** | **Outputs** | **Gap** | **Inputs** | **Outputs** |
| Sensory | 2 | 63 | 1 | 1 | 3 | 3 |
| Sensory/Inter |  |  |  | 1 | 4 |  |
| Interneuron | 2 | 29 | 2 | 4 | 19 | 14 |
| Motor/Inter |  |  | 1 | 3 | 11 | 39 |
| Command/Inter |  |  |  |  |  | 10 |
| Motor |  | 1 |  | 2 |  | 10 |
| Sensory/Motor |  |  |  |  |  |  |

| Quantity and types of synapses Along AIBR Process | | | | | | |
| --- | --- | --- | --- | --- | --- | --- |
| AIBR | **Proximal** | | | **Distal** | | |
|  | **Gap** | **Inputs** | **Outputs** | **Gap** | **Inputs** | **Outputs** |
| Sensory | 3 | 68 | 2 |  | 2 | 1 |
| Sensory/Inter |  |  |  |  | 1 |  |
| Interneuron |  | 24 | 2 | 11 | 43 | 10 |
| Motor/Inter |  |  |  | 3 | 15 | 36 |
| Command/Inter |  |  |  |  | 1 | 9 |
| Motor |  |  |  | 1 | 2 | 14 |
| Sensory/Motor |  |  |  |  | 2 | 1 |

**TABLE 1 LEGEND:** **Composition of AIB Synaptic Partners** Synaptic partner composition of the AIBL (top) and AIBR (bottom) processes. Data obtained from the worm wiring database (<https://wormwiring.org/apps/neuronVolume/>) ^4^. Many neurons were polymodal and were separated accordingly. Numbers here were used to generate nested pie charts in Figure 1A.
